# Intermediate filament dysregulation and astrocytopathy in the human disease model of *KLHL16* mutation in giant axonal neuropathy (GAN)

**DOI:** 10.1101/2023.03.13.532440

**Authors:** Rachel Battaglia, Maryam Faridounnia, Adriana Beltran, Jasmine Robinson, Karina Kinghorn, J. Ashley Ezzell, Diana Bharucha-Goebel, Carsten Bonnemann, Jody E. Hooper, Puneet Opal, Thomas W. Bouldin, Diane Armao, Natasha Snider

**Affiliations:** Department of Cell Biology and Physiology, University of North Carolina at Chapel Hill; Department of Genetics, University of North Carolina at Chapel Hill; National Institute of Neurological Diseases and Stroke, Bethesda, MD; Department of Pathology, Stanford University, Palo Alto, CA; Departments of Neurology and Cell and Developmental Biology, Northwestern University, Chicago, IL; Department of Pathology and Laboratory Medicine, University of North Carolina at Chapel Hill; Department of Radiology, University of North Carolina at Chapel Hill

## Abstract

Giant Axonal Neuropathy (GAN) is a pediatric neurodegenerative disease caused by *KLHL16* mutations. *KLHL16* encodes gigaxonin, a regulator of intermediate filament (IF) protein turnover. Previous neuropathological studies and our own examination of postmortem GAN brain tissue in the current study revealed astrocyte involvement in GAN. To study the underlying mechanisms, we reprogrammed skin fibroblasts from seven GAN patients carrying different *KLHL16* mutations to iPSCs. Isogenic controls with restored IF phenotypes were derived via CRISPR/Cas9 editing of one patient carrying a homozygous missense mutation (G332R). Neural progenitor cells (NPCs), astrocytes, and brain organoids were generated through directed differentiation. All GAN iPSC lines were deficient for gigaxonin, which was restored in the isogenic control. GAN iPSCs displayed patient-specific increased vimentin expression, while GAN NPCs had decreased nestin expression compared to isogenic control. The most striking phenotypes were observed in GAN iPSC-astrocytes and brain organoids, which exhibited dense perinuclear IF accumulations and abnormal nuclear morphology. GAN patient cells with large perinuclear vimentin aggregates accumulated nuclear *KLHL16* mRNA. In over-expression studies, GFAP oligomerization and perinuclear aggregation were potentiated in the presence of vimentin. As an early effector of *KLHL16* mutations, vimentin may serve as a potential therapeutic target in GAN.

## Introduction

Giant Axonal Neuropathy is a rare, progressive, pediatric neurodegenerative disease that affects both the peripheral nervous system (PNS) and the central nervous system (CNS) (1–3). In children affected by GAN, early milestone development is normal (4). Disease onset is usually 3-4 years of age and is often heralded by clumsiness of gait. By the end of the second decade of life, most patients are non-ambulatory and have limited use of their arms and little to no use of their legs (1). Death nearly always occurs by the third decade. The clinical findings and pathologic distribution of lesions in GAN are consistent with a central-peripheral distal axonopathy (5, 6).

GAN is caused by recessive loss-of-function mutations in the *KLHL16* gene (also known as *GAN*) that encodes gigaxonin, an E3 ubiquitin ligase adaptor protein (7). Gigaxonin promotes degradation of many members of the intermediate filament (IF) gene family, which are cell type-specific cytoskeleton components (8–10). In the absence of functional gigaxonin, IF proteins accumulate in many cell types, including melanocytes, endothelial cells, lens epithelial cells, Schwann cells, astrocytes and neurons (11–14). In 1972, the first reported case of GAN described a six-year-old child with an unusual, slowly progressive, sensorimotor neuropathy, and the details of the nerve biopsy**—**fiber loss and “*masses of neurofilaments distending many of the axons to giant proportions*” **—**christened the disease (15). Since that time, validation of therapeutic efficacy and viral vector delivery systems in GAN knock out (KO) rodent models (16–18) have provided the springboard for a first-in-human phase I clinical trial of intrathecal gene transfer of *KLHL16*. To date, most attention has been directed at IF aggregates within GAN neurons and far less attention has been directed at IF aggregates within GAN astrocytes. Despite the fact that conspicuous and abundant IF aggregates within astrocytes are described in every reported autopsy case of GAN (5, 6, 19–21), the contribution of astrocyte dysfunction to neurodegeneration and the pathogenesis of astrocyte IF aggregation remain underinvestigated.

In this work, we use patient-derived induced pluripotent stem cells (iPSCs) and directed differentiation to develop human astrocyte and brain organoid models of GAN. These models reproduce astrocyte IF protein aggregation comparable to astrocytes of GAN patients *in vivo*. The GFAP IF protein inclusions observed in GAN iPSC-astrocytes are accompanied by nuclear invaginations, similar to those recently reported in Alexander Disease (AxD) iPSC-astrocytes (22), suggesting possible shared mechanisms. Additionally, we reveal new mechanisms regarding the role of vimentin in GFAP aggregation and *KLHL16* localization in cells.

## Results

### Astrocytopathy in human GAN

Postmortem histologic examination of cerebral white matter in a GAN patient revealed extensive astrocyte involvement with the presence of innumerable RFs and reactive astrocytosis (**Fig.1A-B**). RFs were concentrated in astrocyte endfeet around blood vessels **(Fig.1A)** and dispersed in cerebellar white matter in proximity to giant axonal swellings **(Fig.1B)**. These data clearly demonstrate that *KLHL16* loss-of-function mutation leads to astrocytes acquiring marked histopathologic features, in agreement with early neuropathologic observations in GAN (5, 6, 19–21). In order to examine the mechanistic link between *KLHL16*/gigaxonin downregulation and/or dysfunction and astrocytopathy, we reprogrammed seven skin fibroblast lines from GAN patients with different *KLHL16* mutations **(Fig.1C; Supplemental Fig.1)** to iPSCs. The GAN cell lines contained unique *KLHL16* mutations that span all functional domains of gigaxonin (23), including null mutations (patients 3, 4), mutations in the BTB (patient 2), Back (patient 3), and Kelch (patients 1, 5, 6, 7) domains **(Fig.1D)**. Isogenic controls of the two lines carrying homozygous missense mutations (patients 2 and 7) were generated by CRISPR/Cas9 gene editing **(Supplemental Fig.2A-F)**. However, despite the correction of the known mutation in Patient 2 (Y89S), IF protein aggregation persisted in the isogenic control iPSC-astrocytes **(Supplemental Fig.2G-H)**, suggesting the potential involvement of other mutations that have yet to be identified in this patient. For these reasons, the majority of this work focused on Patient 7 carrying the G332R Kelch-domain mutation and corresponding isogenic clones that we generated from the parental line. Because the Kelch domain mediates the associations between gigaxonin and IF proteins (24, 25), we specifically focused on IF phenotypes in iPSC, NPCs, astrocytes and brain organoids **(Fig. 1E)** generated from patient 7 cells and isogenic controls.

**Figure 1.**
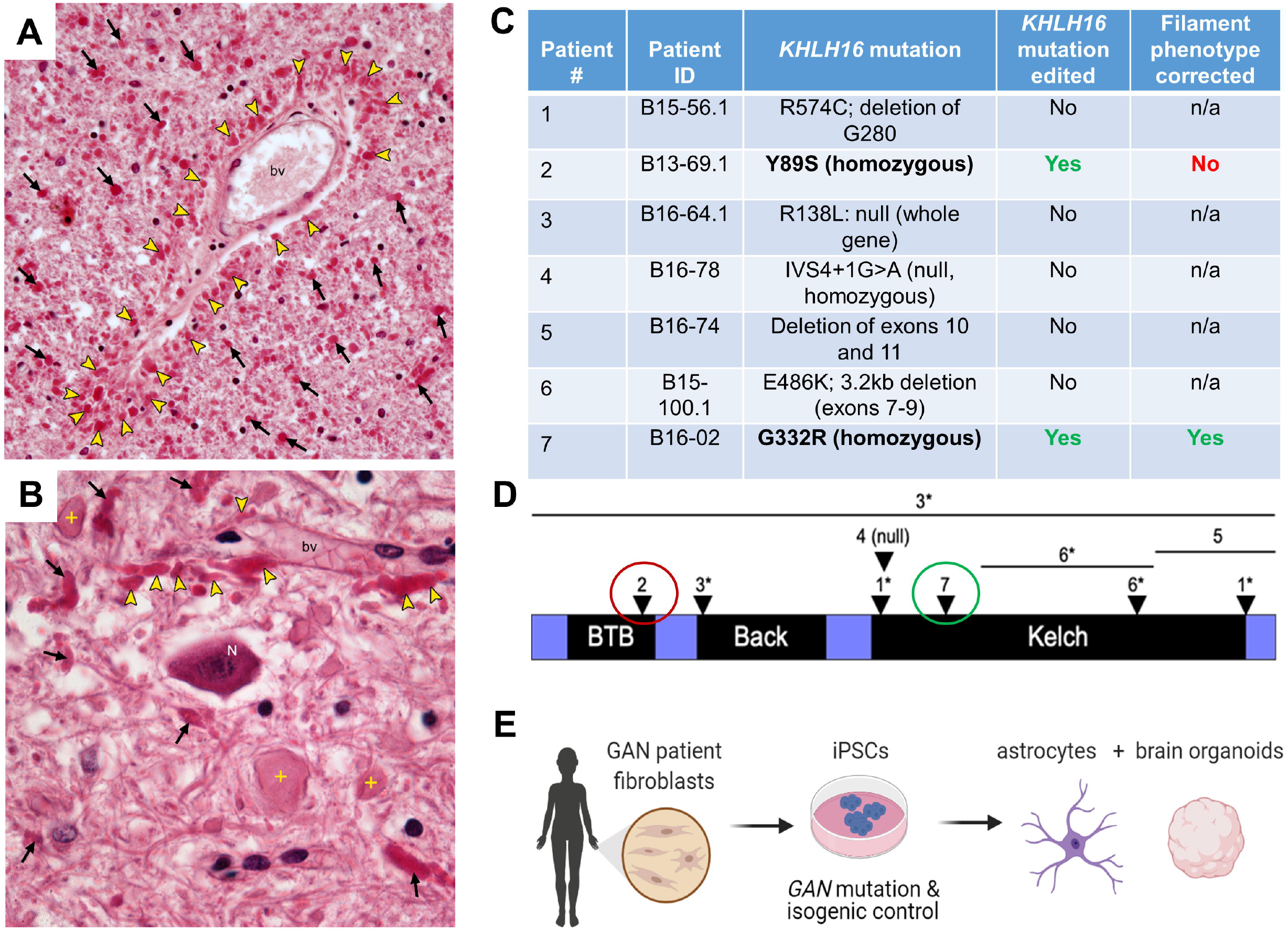
Modeling GAN-related astrocytopathy using patient-derived iPSCs. **(A)** Abundant, bright red, oval and club-shaped astrocyte inclusions, or Rosenthal fibers (RFs; arrowheads) clustered around a blood vessel (bv) in GAN. Note innumerable RFs (arrows) and pathologic remodeling of astrocytes in cerebral white matter (H&E;40X). **(B)** RFs are concentrated in astrocyte endfeet (arrowheads) around a blood vessel (bv). RFs and giant axonal swellings (crosses) are dispersed in cerebellar white matter surrounding the dentate nucleus. A neuronal cell body (N) is morphologically unremarkable (H&E; 60X). Decedent was a young child with phenotypically classic GAN. **(C)** Summary of GAN patient cell lines with their known mutations that were used in the study. The two lines carrying homozygous missense mutations (Y89S and G332R) were edited to generate isogenic clones. Only one (patient 7) of the two isogenic lines displayed normal filaments upon differentiation and was used in subsequent experiments. **(D)** Schematic of human *KLHL16* mutations mapped to the protein, gigaxonin. Numbers 1-7 refer to the patient donor cells that were used in this study; listed in Panel C. Asterisks indicate compound heterozygous mutations. Red circle indicates isogenic line was generated, but retained abnormal IF phenotype. Green circle indicated isogenic line was generated and it resulted restored IF phenotype. Please also see Supplemental Figure 2. **(E)** Schematic diagram of methods used to generate isogenic control and GAN iPSC-astrocytes and brain organoids from Patient 7 (G332R).

### Patient-specific upregulation and perinuclear bundling of vimentin in GAN iPSCs

We observed by immunoblot analysis that gigaxonin protein level was largely undetectable or extremely low in all of the seven GAN iPSC lines when compared to a non-GAN control iPSC line (15CA) as a reference (26). Gigaxonin expression in the four isogenic control lines of patient 7 (B3,G5,2D1, and 2D3) was restored to levels similar of those in the 15CA non-GAN control **(Fig.2A)**. We did not observe major differences in lamin-A/C and B1 expression **(Fig.2A)** or Lamin B1 localization **(Supplemental Fig.3)** in the parental line versus the isogenic control cells. However, by immunoblot we observed variable vimentin expression, such that four patient lines (2, 5, 6, 7) expressed detectable levels of vimentin (Vim^+^), while the remaining patient lines (1,3,4) did not (Vim^-^), similar to the 15CA non-GAN control and the patient 7 isogenic clones **(Fig. 2A)**. Immunofluorescence analysis of the patient 7 (G332R) iPSCs and two of its isogenic clones (2D1 and 2D3) also showed that vimentin expression **(Fig.2B-C)** was significantly higher in iPSCs colonies of the mutant line compared to the isogenic controls (p<0.05; one-way ANOVA). Keratin 8 filament staining was used as another cytoplasmic IF marker of the iPSC colonies **(Fig.2B)** and appeared filamentous across lines, although detailed ultrastructural analysis and quantitative examination of the keratin network dynamics were not performed as part of this study. Differences in vimentin gene (*VIM*) expression did not account for the protein variation among patient lines **(Fig. 2D)**. We also tested by qPCR the expression of all *KLHL* genes using sequence-specific primers **(Supplemental Table 3)** and observed that the expression of five *KLHL* genes (*KLHL*-12/19/36/39/42) was significantly decreased in the Vim^+^ compared to the Vim^-^ iPSCs. These results suggest that the phenotypic variability in vimentin expression in the setting of *KLHL16* mutations may potentially be influenced by the expression of related *KLHL* genes.

**Figure 2.**
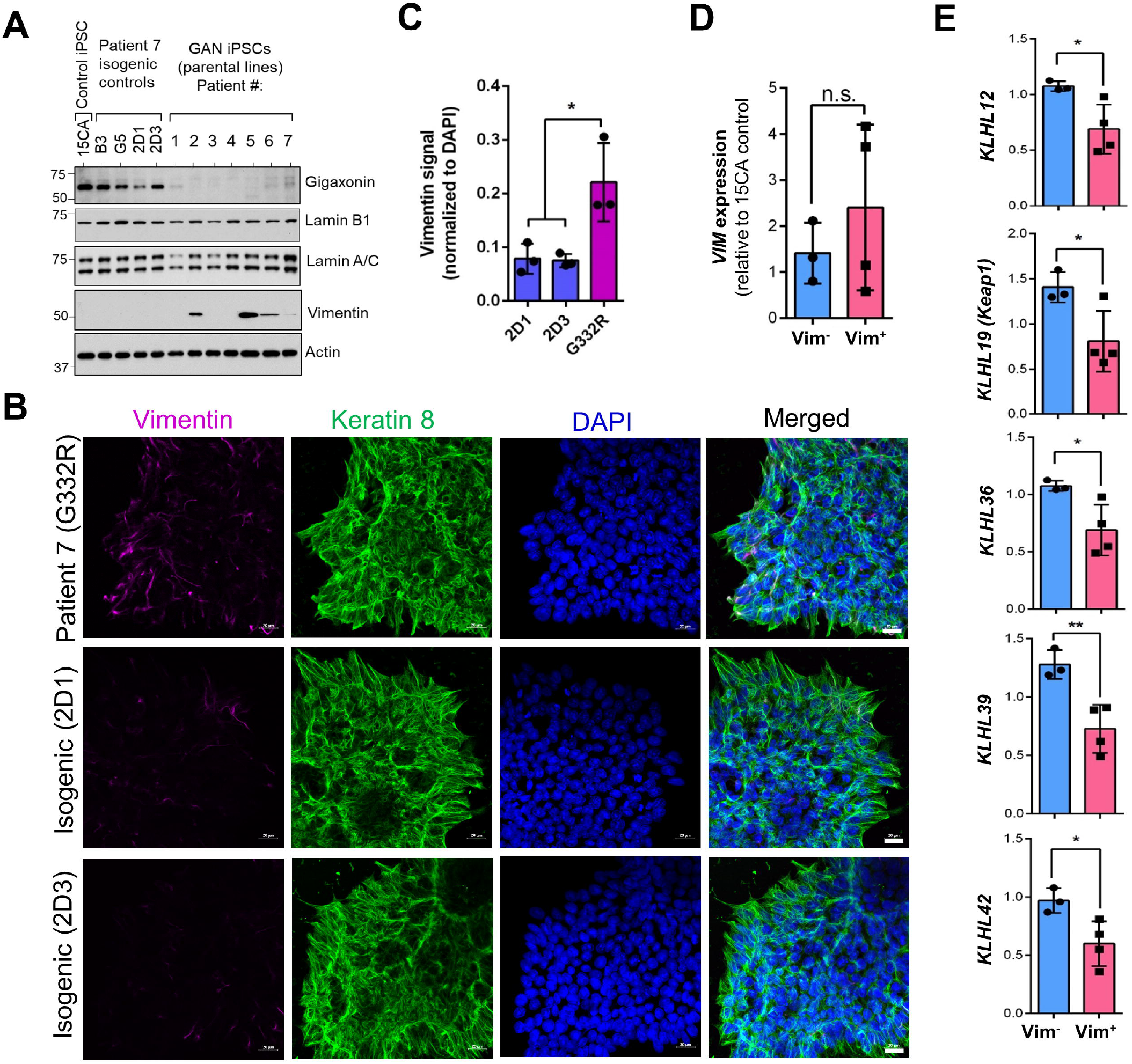
Human *KLHL16* loss-of-function mutations selectively alter vimentin expression in iPSCs. **(A)** Immunoblotting for gigaxonin and IF proteins lamin B1, lamin-A/C and vimentin in total lysates from non-GAN control (15CA), four isogenic iPSC controls of GAN Patient #7 (B3,G5,2D1,2D3) and seven unedited GAN iPSCs parental lines (patients 1-7). Actin blot serves as protein loading control. **(B)** Immunofluorescence analysis of vimentin (magenta), Keratin 8 (green) and DAPI (blue) in patient 7 (G332R) and isogenic control iPSC clones 2D1 and 2D3 from the same patient. Scale bars=20_μ_m. **(C)** Vimentin immunofluorescence intensity quantified by Image J. n=3 fields of view per condition (equal 1228800 pixel areas measured and normalized to DAPI signal) *p<0.05; One-way ANOVA. **(D)** Quantitative real-time PCR analysis of *VIM* gene expression in Vim^-^ (Patients 1,3,4) vs Vim^+^ (Patients 2,5,6,7) GAN iPSCs. n.s.=not significant; unpaired t-test. **(E)** Quantitative real-time PCR analysis of *KLHL* gene expression in Vim^-^ (n=3) vs Vim^+^ (n=4) GAN iPSCs. *p<0.05; **p<0.01; unpaired t-test.

### Reduced nestin expression in GAN neural progenitor cells (NPCs) compared to isogenic controls

Next, we differentiated the iPSCs to neural progenitor cells (NPCs), which express the type IV IF protein nestin. Immunoblot analysis of IF proteins revealed similar expression of lamin-A/C and vimentin, but 3-fold higher (p<0.05; two-way ANOVA) nestin expression in the isogenic controls compared to the parental line G332R **(Fig.3A-B)**, which was corroborated by immunofluorescence analysis **(Fig.3C)**. These data suggest GAN iPSCs may have reduced *in vitro* potential to differentiate to nestin-positive NPCs, which may be linked to previously reported abnormality in the Sonic hedgehog (Shh) pathway in GAN (27).

**Figure 3.**
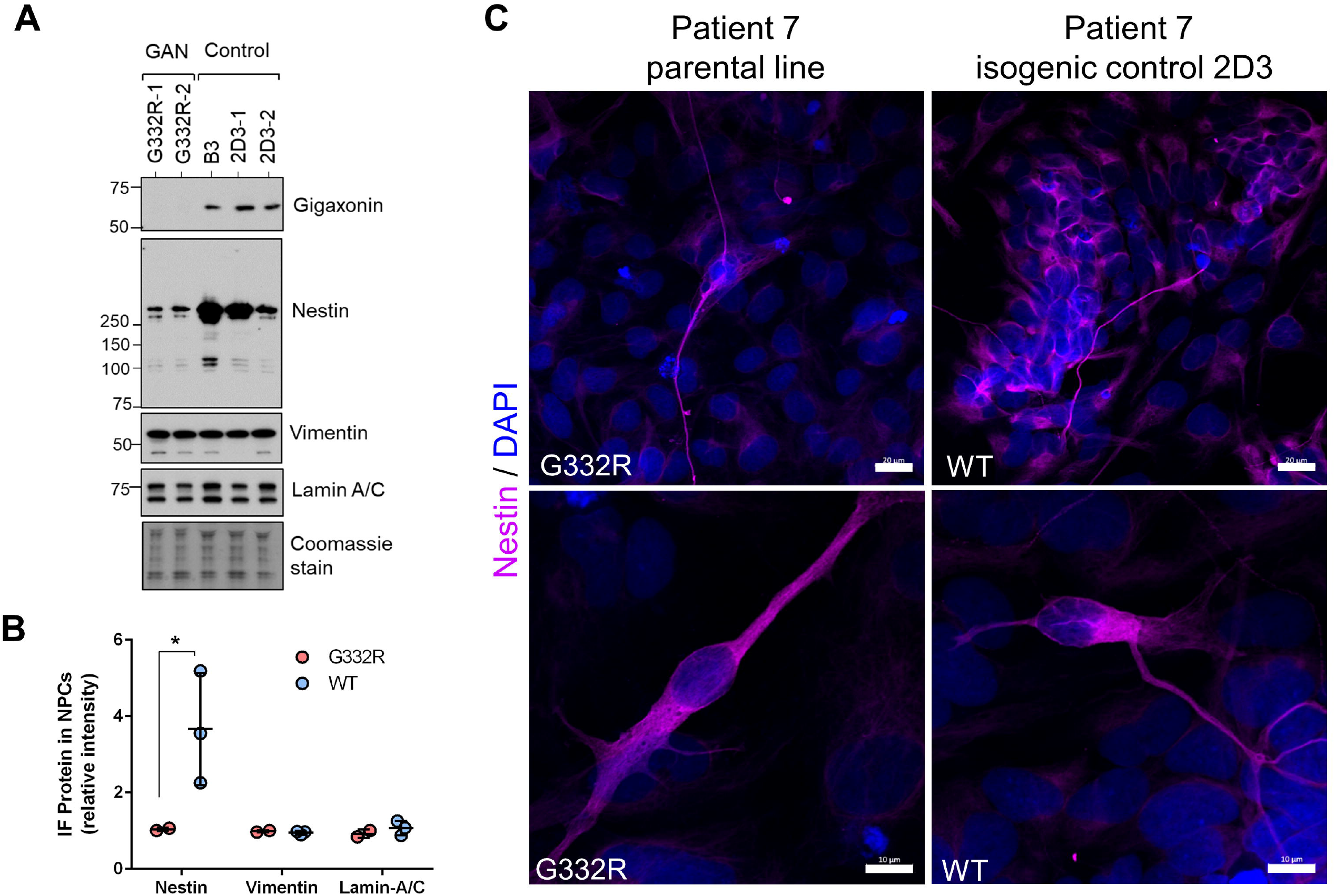
Reduced nestin expression in GAN mutant neural progenitor cells (NPCs) compared to isogenic controls. **(A)** Immunoblotting for gigaxonin, nestin, vimentin, and lamin A/C in GAN (n=2) and isogenic (n=3) NPCs derived from Patient #7 (G332R). Coomassie stain serves as total protein loading control. **(B)** Quantification of band intensities for G332R (n=2) and control (n=3) from panel A. *p<0.05 two-way ANOVA. **(C)** Confocal images of immunofluorescence of NPCs stained for nestin (magenta) and DAPI (blue). Scale bars=20 μm (top) and 10 μm (bottom).

### Perinuclear accumulations of vimentin and GFAP in GAN iPSC-astrocytes

Given the observed early progenitor phenotypes, we next asked if GAN NPCs could be differentiated to astrocytes in culture. Using defined differentiation media (22), we were able to generate both GAN mutant (G332R) and corresponding isogenic control vimentin-positive astrocytes at day 60 of the differentiation procedure (**Fig. 4A)**. One major difference between control and GAN iPSC-astrocytes at this stage was the presence of large, intensely stained, perinuclear vimentin accumulations in the latter **(Fig. 4B-C)** that was not due to cell death or overall change in cell size **(Supplemental Fig 4)**. Consistent with this perinuclear concentration, the vimentin-occupied areas were significantly smaller in the GAN mutant cells **(Fig. 4D)**. At this time point (60 days) ALDHL1 expressing cells comprised approximately 12% and 10% in the isogenic and mutant cells, respectively **(Supplemental Figure 5)** while GFAP-positive cells comprised approximately 10% and 7% in the isogenic and mutant cells, respectively, compared to the total number of cells **(Figure 4E-F)**. The GFAP-positive GAN iPSC-astrocytes contained large perinuclear GFAP aggregates and displayed misshapen nuclei along with weak and diffuse abnormal cytoplasmic lamin B1 staining **(Fig.4G)**. This phenotype was in stark contrast to the isogenic control astrocytes, which displayed normal GFAP IF organization, normal nuclear morphology, and normal lamin B1 staining **(Fig. 4G)**. In addition to vimentin, GFAP-positive inclusions in GAN iPSC-astrocytes stained positive for other RF markers, including ubiquitin, _α_B-crystallin, and Hsp27 **(Supplemental Fig.6)**. Collectively, our data show that GFAP and vimentin co-accumulate in the perinuclear region of GAN iPSC-astrocytes, and that these accumulations are associated with abnormalities of nuclear morphology. These observations were very similar to our recent findings in AxD iPSC-astrocytes, where GFAP aggregates are observed adjacent to highly dysmorphic nuclei (22).

**Figure 4.**
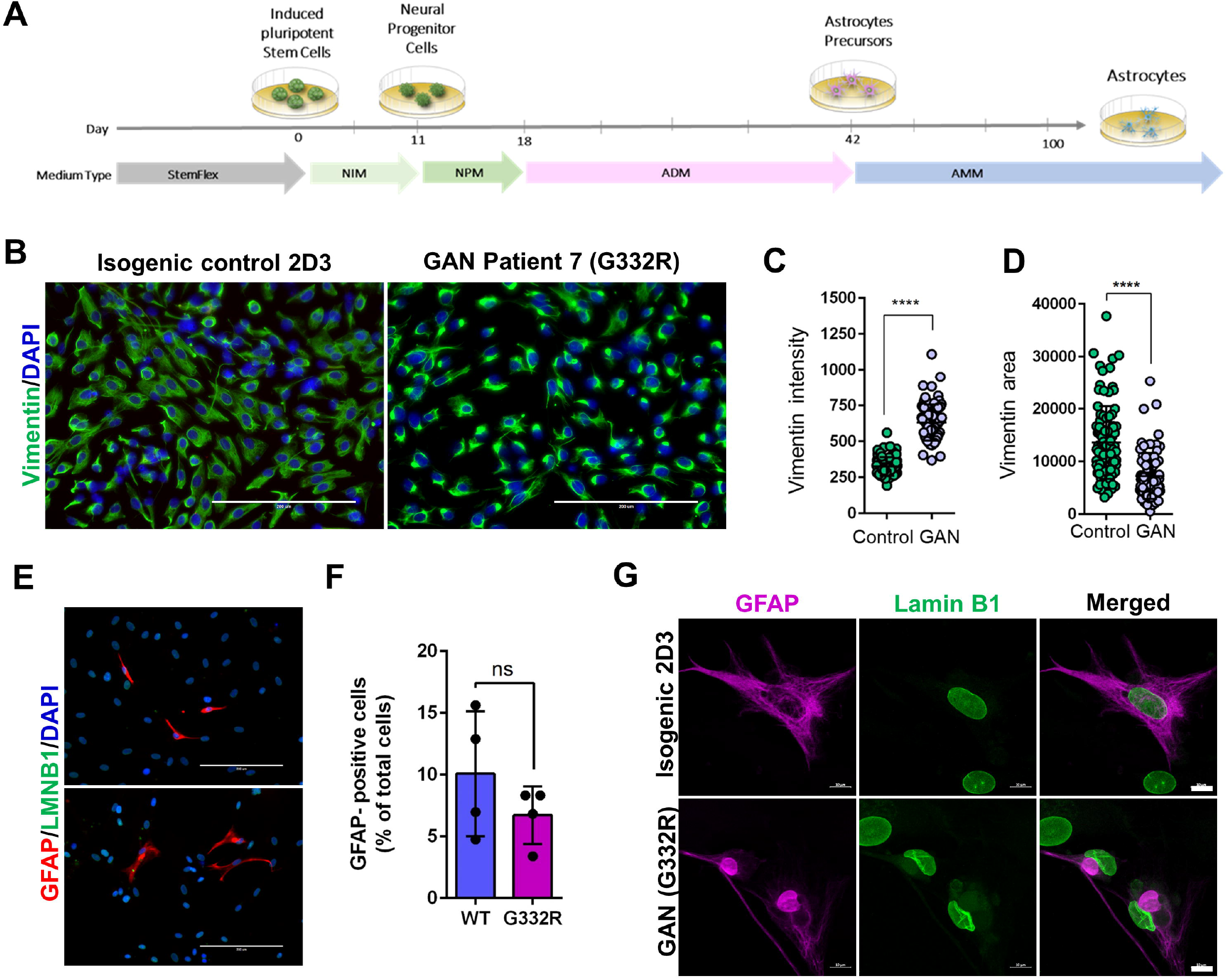
GAN iPSC-astrocytes display perinuclear vimentin and GFAP aggregates and abnormal nuclei. **(A)** Schematic illustrating the workflow for iPSC differentiation to astrocytes. **(B)** Vimentin bundling in immature GAN iPSC-astrocytes analyzed at day 60 of the differentiation timeline. Scale bars=200μm. **(C)** Quantification of the vimentin signal intensity in control and GAN iPSC-astrocytes. ****p<0.0001; unpaired t-test. **(D)** Quantification of vimentin area in control and GAN astrocytes. p<0.0001; unpaired t-test. **(E)** GAN patient 7 (G332R) and corresponding isogenic control (clone 2D3) iPSC-astrocytes were stained for GFAP (red), lamin B1 (green), and DAPI (blue) Scale bars=200 μm. **(F)** Quantification of GFAP-positive cells as a percentage of total cells from 4 representative fields of view (n=148 WT; n=199 G332R; total number of cells counted). **(G)** GAN and isogenic control iPSC-astrocytes were stained for GFAP (magenta) and lamin B1 (green). Scale bars=10 μm.

### GAN brain organoids express GFAP and neurofilaments and exhibit cytoskeletal and nuclear abnormalities

Because few iPSC-astrocytes expressed GFAP in the monoculture system, we sought a new model to generate greater numbers of GFAP-positive astrocytes. To that end, we differentiated iPSCs to brain organoids using an established protocol (**Figure 5A**) (28). After 7 months of *in vitro* culture, the isogenic (2D3 clone) and the corresponding mutant (G332R mutation) GAN brain organoids were enriched in GFAP-positive astrocytes that were integrated with other cell types, including neurofilament-M/H expressing neurons **(Fig. 5B)**. At the ultrastructural level, cellular morphology appeared normal in isogenic organoids, reflected in evenly spaced IFs running in parallel to one another and unremarkable organelles, like mitochondria **(Fig. 6; top panels)**. In contrast, GAN organoids displayed tightly bundled, swirling, abnormal IF accumulations that trapped smaller organelles, like mitochondria **(Fig. 6; bottom panels)**. Furthermore, and in line with the phenotype of the iPSC-astrocyte monoculture system, the organoids contained abnormal IF accumulations encroaching upon irregularly contoured nuclei displaying compromised integrity of the nuclear envelope. Although the ultrastructural examination of the organoids reveals GAN phenotypes that are observed *in vivo*, our present analysis did not distinguish between neurons and astrocytes, as both are affected in GAN cells.

**Figure 5.**
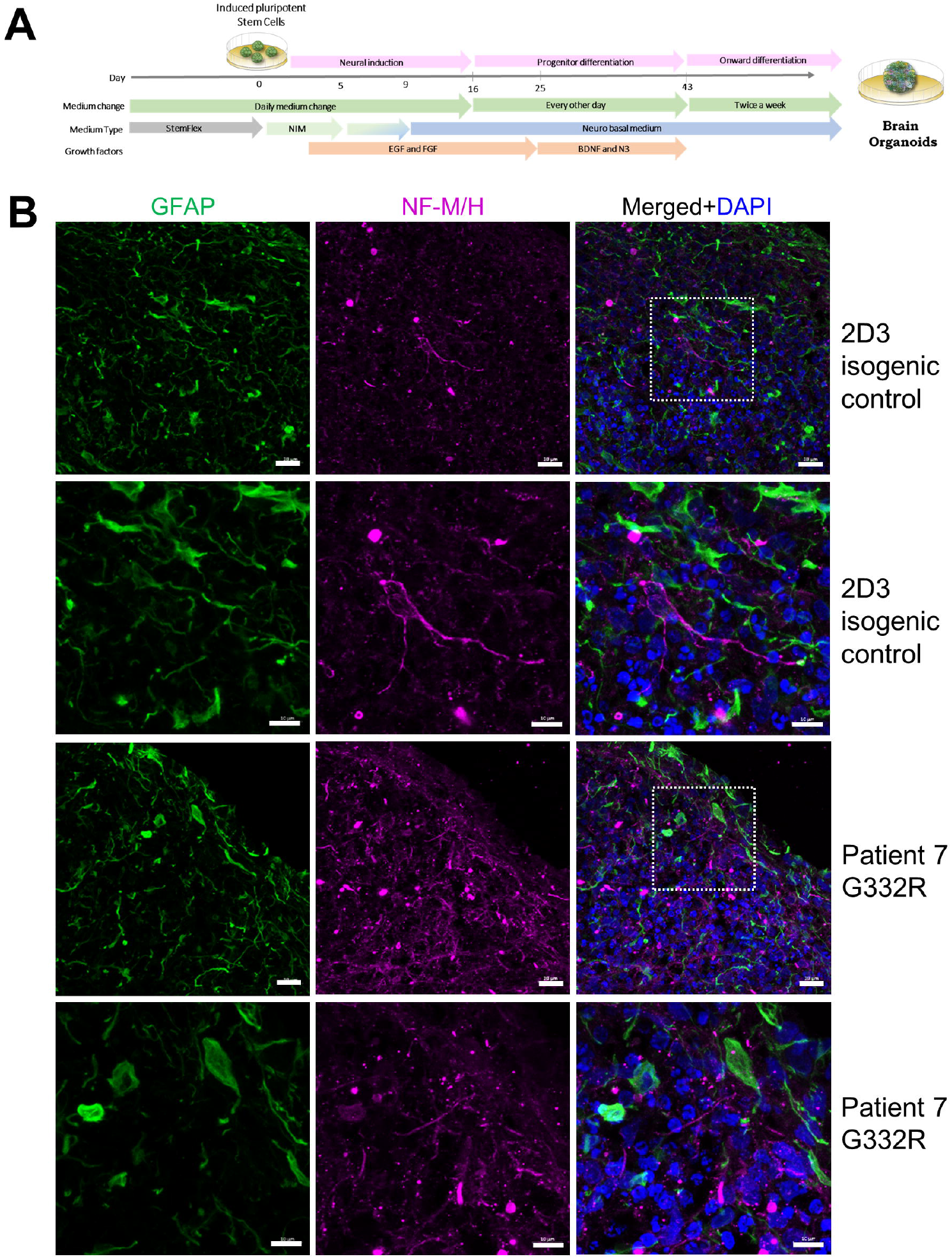
Generation of GAN and isogenic control iPSC-derived brain organoids **(A)** Schematic diagram of methods for generating brain organoids from iPSCs. **(B)** Confocal images of immunofluorescence staining for GFAP (green), neurofilaments-M/H (magenta) and DAPI (blue) in GAN patient 7 (G332R) and corresponding isogenic control brain organoid clone 2D3. Scale bars=20_μ_m (top and third row) and 10_μ_m (second and fourth row).

**Figure 6.**
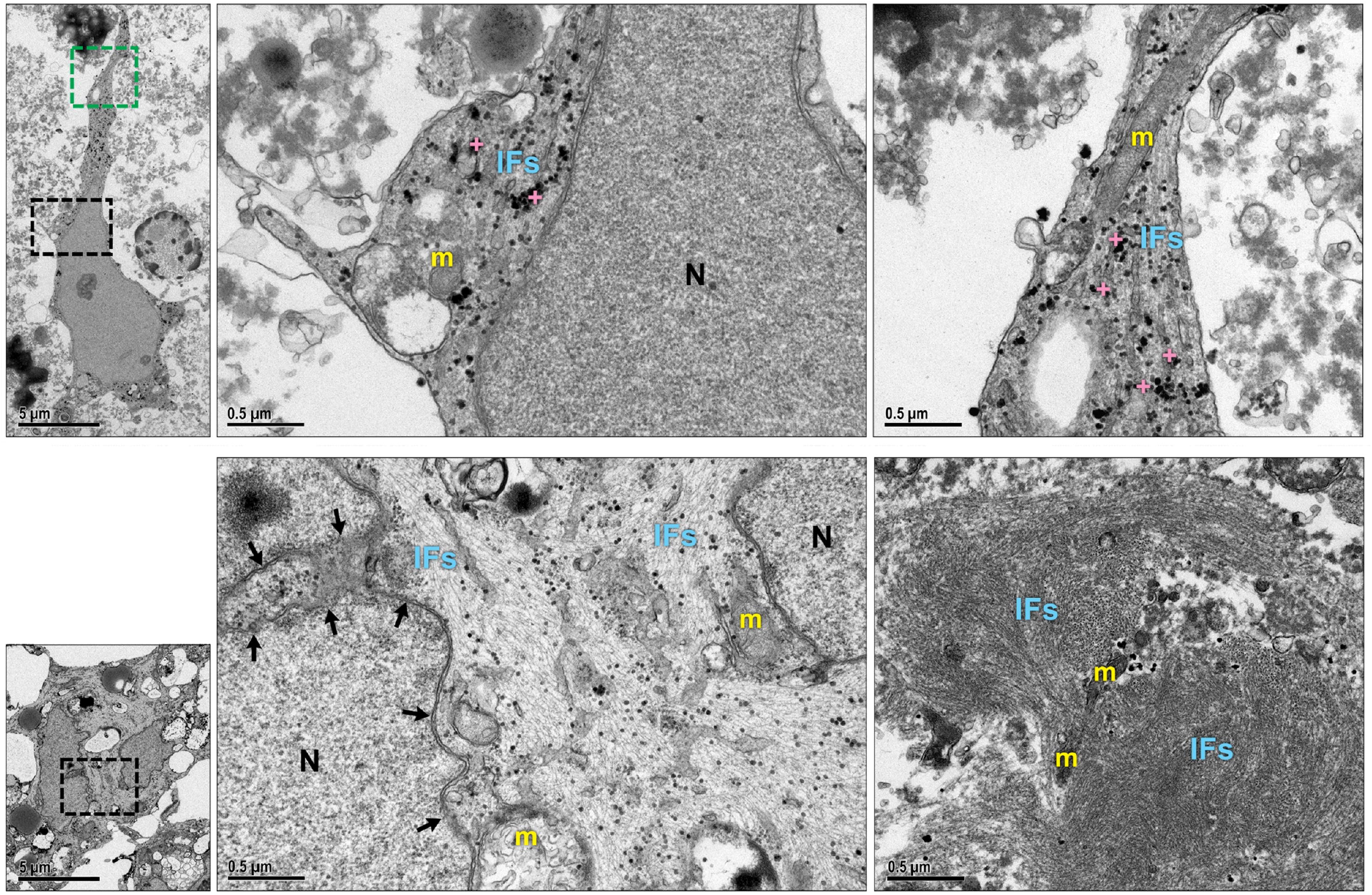
GAN brain organoids recapitulate IF dysregulation and nuclear abnormalities. **Top** left panel shows EM image of an isogenic control brain organoid revealing a star-shaped cell with a regularly contoured nucleus and multiple, slender cytoplasmic processes.[5kx; scale bar= 5 _μ_m]. Top middle panel shows enlargement of black boxed area. Note abundant glycogen granules (crosses), ultrastructurally unremarkable mitochondria (m), smoothly contoured nucleus (N), and evenly dispersed, parallel arrays of intermediate filaments (IFs)[50 kx; scale bar= 0.5 _μ_m]. Top right panel shows enlargement of green boxed area. Note unremarkable mitochondria (m), glycogen granules (crosses) and normally distributed and aligned intermediate filaments (IFs [50 kx; scale bar=0.5 _μ_m] **Bottom** left panel shows EM image of a GAN brain organoid revealing a cell with an irregularly contoured nucleus. [5kx; scale bar= 5 _μ_m]. Bottom middle panel shows enlargement of boxed area. Note abnormal intermediate filament accumulations (IFs) encroaching upon an irregularly contoured nuclear envelope (arrows) and enmeshed mitochondria (m). [50kx; scale bar= 0.5 _μ_m]. Bottom right panel shows a GAN brain organoid revealing tightly bundled, swirling abnormal IF accumulations and IF-entrapped organelles, including mitochondria (m). [50kx; scale bar= 0.5 _μ_m]

### Increased perinuclear GFAP accumulation in the presence of vimentin IFs

To probe the role of vimentin in GAN astrocytes, we utilized an assembly-compromised and aggregation-prone S13D-GFAP phosphomimic mutant relevant to AxD astrocytes, which we characterized previously (22). As detailed previously (22), we used SW13 cells specifically because the Vim+ SW13 sub-clone of this cell line expresses only vimentin IFs (and no other cytoplasmic IFs), while the vim-SW13 sub-clone lacks all cytoplasmic IFs (29, 30), making these cells a suitable transfection host to determine how vimentin affects GFAP organization. We observed that the perinuclear aggregation of S13D-GFAP was augmented in the presence of vimentin **(Fig. 7A)**. Notably, smaller spherical S13D-GFAP aggregates appeared to associate with the vimentin network in and around the perinuclear region **(Fig. 7B)**. In addition, we observed 10-fold elevated levels of high-molecular-mass S13D-GFAP oligomers in vimentin-positive compared to vimentin-negative SW13 cells **(Fig.7C-D)**. These results suggest that the high level of expression and perinuclear accumulation of vimentin may be an important driver of astrocyte pathology in GAN, in part by facilitating GFAP aggregation.

**Figure 7.**
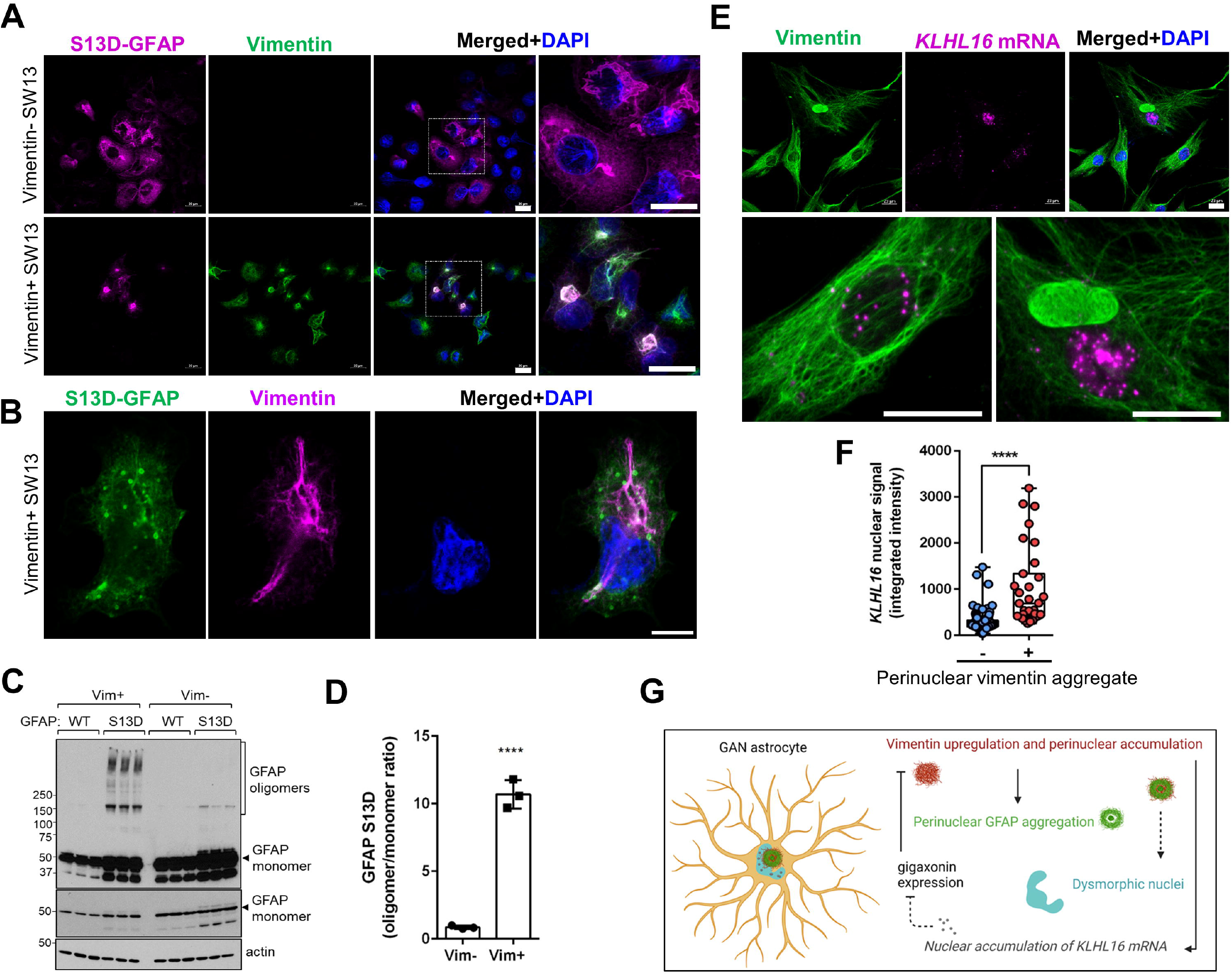
Increased GFAP aggregation and nuclear *KLHL16* mRNA accumulation in the presence of vimentin IFs and perinuclear aggregates. **(A)** Confocal images of immunofluorescence staining for GFAP (magenta), vimentin (green), and DAPI (blue) in 24h-transfected Vim- (top) and Vim+ (bottom) SW13 cells. Scale bars=20μm. Images on the far right show the selected enlarged areas. **(B)** Immunofluorescence analysis of S13D-GFAP (green), vimentin (magenta) and DAPI (DNA; blue) after 12h of transfection. Note the co-localization of the smaller spherical GFAP aggregates along vimentin filaments. **(C)** Immunoblot analysis of GFAP and actin (loading control) of lysates from SW13 vimentin-positive and vimentin-negative SW13 cells. Note the abundance of S13D-GFAP oligomers in the presence, but not in the absence, of vimentin. Middle panel is a lighter exposure of the top membrane to show the GFAP monomer. **(D)** Densitometry quantification of the GFAP oligomer/monomer ratios from the panel B. *p<0.0001; unpaired t-test. **(E)** Co-staining of vimentin (green) and *KLHL16* mRNA encoding gigaxonin (magenta) in GAN patient fibroblasts. Note the significant nuclear accumulation of *KLHL16* mRNA in the vimentin aggregate-containing cell compared to the surrounding cells without aggregates. Scale bars=20μm. Bottom, magnified images of *KLHL16* mRNA (magenta) in a cell without (left) and with (right) vimentin (green) aggregate. **(F)** Quantification of *KLHL16* mRNA nuclear signal intensity in GAN fibroblasts with (n=31) and without (n=143) perinuclear vimentin aggregates. *p<0.0001; unpaired t-test. **(G)** Working hypothesis for proteostasis failure in GAN astrocytes. Vimentin exacerbates astrocyte pathology by promoting GFAP aggregation and nuclear accumulation of *KLHL16* mRNA, which in turn aggravates further IF accumulation due to lack of gigaxonin protein translation.

### Nuclear enrichment of KLHL16 mRNA in the presence of perinuclear vimentin aggregates

Given the deficiency of gigaxonin protein in the GAN patient cells, irrespective of the mutation (Fig.2A), we asked whether the *KLHL16* mRNA is sequestered inside IF aggregates. To address this question, we analyzed GAN patient fibroblasts, which can express normal vimentin IFs alone, or together with large perinuclear aggregates (25). Although we did not observe co-localization between *KLHL16* and vimentin aggregates, there was a strong nuclear presence of *KLHL16* in the cells with large perinuclear aggregates compared to cells without such inclusions **(Fig. 7E-F)**. This raises the possibility that IF aggregation may impede the nuclear transport and translation of *KLHL16* to gigaxonin, further exacerbating progressive IF accumulation and cellular dysfunction **(Fig.7G)**. This remains to be rigorously tested. Our results here demonstrate that the morphologic perturbations induced by IF protein accumulation may have functional consequences on the aberrant processing of *KLHL16* mRNA in GAN patient cells. These observations highlight the importance of human-specific disease models of GAN, such as the one described herein, to uncover disease mechanisms and identify molecular targets for therapy.

## Discussion

In this study, we developed a model to investigate IF proteostasis failure in human astrocytes from GAN patients. Utilizing GAN patient-derived iPSCs and genetically engineered isogenic controls of a single patient line (G332R), we showed that *KLHL16* mutations depleted gigaxonin protein and compromised vimentin and nestin expression in iPSCs and NPCs, respectively. We also detected similarities between GAN and AxD iPSC-astrocytes, specifically the presence of nuclear abnormalities associated with cytoplasmic GFAP+/Vim+ aggregates. We extended this work to 3D iPSC-brain organoids that reflected IF pathology that is observed *in vivo* in GAN patients. Finally, we showed increased oligomerization and aggregation of assembly-compromised GFAP and accumulation of nuclear *KLHL16* in the presence of vimentin. Thus, our findings confirm that GAN iPSC-astrocytes and brain organoids are a valuable experimental tool to examine uniquely human disease mechanisms in GAN **-** mechanisms that implicate vimentin as an underappreciated contributor to the astrocytopathy observed in GAN.

Our model expands upon a previous human iPSC-neuron model of GAN by directing attention to the contributions of astrocytes (31). The human iPSC-astrocyte and organoid models used in this work offer several advantages in the study of GAN and neurologic diseases associated with pathologic remodeling of astrocytes and neurodegeneration. For example, iPSC-derived patient astrocytes show potential for expressing disease-relevant phenotypes, accelerating characterization of pathologic pathways and providing a tractable platform for development of astrocyte-specific drug therapy. Human astrocytes differ from their rodent counterparts in size, shape, and function (32). To examine astrocyte-related disease mechanisms, it is necessary to complement animal models with a ‘humanized’ model faithful to mirroring distinctly human properties. Advances in directed differentiation methods in combination with iPSC technology provide a minimally invasive option to generate human astrocytes (28, 33). Further, organoid-derived astrocytes have demonstrated several astrocyte functions, and thus, this model can be utilized in the future to determine astrocyte functional deficits in GAN (34). The use of patient-derived cells also allows for tailoring of background genetics. This is especially helpful in studies of the role of astrocytopathy in GAN, where there is phenotypic variability in disease severity likely due to genetic modifiers that may not be reflected in animal models.

Species differences in the functional and pathologic phenotype of GAN are apparent when comparing humans to canines, rodents, and zebrafish. From both a clinical and pathologic standpoint, canine GAN most closely mirrors human GAN (35). Canine GAN disease onset is early in life (∼15 months), heralded by hindlimb signs including dragging of toes, diminished proprioception, progressive ataxia and paresis (36–38). In canine GAN, death is usually by 2-years-of-age, and the clinical and pathologic phenotype is compatible with a central-peripheral distal axonopathy (35, 39). In contrast, while rodent models demonstrate similar human GAN pathology, they have normal lifespans and only mild motor and sensory deficits that are late onset (40–42). The zebrafish model of GAN has locomotion defects but also displays developmental defects in motor neurons that are not seen in other models, making the motor phenotype difficult to compare to other species (43). The availability of human cellular disease models of GAN can help bridge the gap in our understanding of disease mechanisms based on other species. However, we acknowledge that disparities in GAN severity across species are multifactorial, including lifespan, length differences in axons, and differing complexity of cells (e.g., astrocyte complexity), such that the various aspects of cell biology, including astrocyte homeostasis and astrocyte-neuron interactions must be tested experimentally.

The recessive nature of GAN and the absence of detectable gigaxonin protein has led to the understanding that the disorder is caused primarily by loss of function of gigaxonin. In the case of null mutations, the cause of gigaxonin depletion is clear, but for missense mutations, it is not fully understood why gigaxonin protein is lost. It has been proposed that the defect occurs post-translationally because missense mutations could destabilize the gigaxonin protein (44). However, this hypothesis is yet to be tested experimentally, and we suggest that a key intermediate step—translation—should be examined. It is plausible that *KLHL16* is transcribed and stalled at the mRNA transcript stage, an idea that is supported by the finding that *KLHL16* mRNA can be stored in stress granules in human cells (45). The possibility that posttranscriptional mechanisms may contribute to GAN raises the question of whether GAN can be caused by mutations outside the *KLHL16* coding region. Although some mutations have been identified in splicing sites, sequencing is not routinely performed on non-coding regions of *KLHL16* in suspected GAN patients, which is understandable considering the remarkable length of the *KLHL16* gene (23). This patient-derived iPSC model will provide a useful tool to address the mechanism of loss of gigaxonin in GAN.

Finally, our results indicate vimentin and GFAP IFs are involved in cytoskeletal proteostasis demise in GAN astrocytes. The presence of deep nuclear indentations juxtaposed to cytoplasmic GFAP+ aggregates suggests that GFAP could promote enhanced and possibly toxic interactions between IF aggregates and the nucleus. One possible cause for this could be related to plectin, which is an IF-associated protein that crosslinks IFs to other structures, including microtubules, actin, and the nuclear envelope (46, 47). While both vimentin and GFAP are known to bind to plectin, it is possible that each IF develops distinct post-translational modifications, which can alter plectin binding (46, 48–50). Another potential mechanism could be disrupted nuclear transport. GFAP and vimentin both contain a predicted nuclear localization signal in the C-terminal tail domain, but GFAP harbors an additional predicted domain in the N-terminal head domain (51). However, it has yet to be shown that GFAP is transported into the nucleus, as has been demonstrated with keratin-17, an epithelial IF (51). Using our overexpression system, we also show evidence of enhanced oligomerization and aggregation of assembly-compromised GFAP in the presence of vimentin. We suspect, as others have posited, that vimentin promotes aggregation by providing a scaffold on which aggregates are seeded (52). Astrocytes are not the only cell that express more than one IF in development and disease. In fact, neurons can express at least four IFs: neurofilaments light, medium, and heavy, as well as peripherin or α-internexin, which are assembled into filaments at very specific ratios (53, 54). It is likely that different IFs cooperate to promote aggregation in other IF-associated disorders where IF proteostasis and assembly are compromised. The abundance of IF aggregates in a variety of cell types in GAN makes our iPSC model a clinically relevant and versatile tool to investigate the effects of IF proteostasis failure in diverse human cellular contexts.

Collectively, our results provide a new model to study the impact of *KLHL16* gene mutations to astrocyte cell biology that may contribute to disease pathogenesis in GAN. Future studies will implement this model to characterize the role of vimentin and post-transcriptional mechanisms in GAN-associated astrocytopathy.

## Materials and Methods

### Antibodies

The following primary antibodies and concentrations were used in this study: rabbit anti-GFAP (DAKO, Agilent, clone Z0334; IF 1:500, WB 1:10,000), mouse anti-GFAP (Sigma, clone GA5; IF 1:300), mouse anti-pSer13-GFAP (gift from Dr. Masaki Inagaki, clone KT13, IF 1:20), mouse anti-Gigaxonin (Santa Cruz Biotechnology, F3, WB 1:200), rabbit anti-Vimentin (Cell Signaling Technology, D21H3, IF 1:100), mouse anti-Vimentin (Thermo Fisher Scientific, V9, WB 1:1000), mouse anti-Keratin 8 (Thermo Fisher Scientific, TS1, IF 1:100), rat anti-K8 (Developmental Studies Hybridoma Bank, Troma I, WB 1:5000), rabbit anti-Lamin A/C (Santa Cruz Biotechnology), rabbit anti-Lamin B1 (Abcam ab16048, IF 1:10,000, WB 1:10,000), mouse anti-Nestin (Thermo Fisher Scientific, 10C2, IF 1:200), mouse anti-Tra-1-60 (Thermo Fisher Scientific, 41-1000, IF 1:300), mouse anti-Tra-1-81 (Thermo Fisher Scientific, 41-1100, IF 1:300), rabbit anti-Oct4 (Abcam, ab19857, IF 1:40), and rabbit anti-Sox2 (Thermo Fisher Scientific, 48-1400, IF 1:125). The following secondary antibodies and concentrations were used: Alexa 488- and Alexa 594-conjugated goat anti mouse and rabbit antibodies (Thermo Fisher Scientific, IF 1:500), and peroxidase-conjugated goat anti-mouse and rabbit antibodies (Sigma, WB 1:5000).

### Cell line

SW13vim+ and vim-cells (29, 30) were provided by Dr. Bishr Omary and cultured in DMEM with 10% fetal bovine serum and 1% penicillin-streptomycin. Authentication was done by short tandem repeat (STR) profiling by ATCC. Transfection were performed as described previously (22) and cells analyzed after 24 hours, unless noted otherwise in the Figure Legend.

### Human tissue

Human specimens were obtained through a validated autopsy consent from the legal next of kin, which explicitly stated that tissues could be used for research purposes. The consent and diagnostic autopsy report were filed in the deceased patient’s medical record.

### Cellular reprogramming, characterization, and gene editing of iPSCs

iPSCs were generated from GAN (OMIM#256850) patient fibroblasts and characterized using the methods previously described (55). Fibroblast reprogramming to iPSCs, karyotyping, CRISPR/Cas9 gene editing, screening of edited clones, and off-target analyses were performed as we described in detail previously in studies on AxD iPSC-astrocytes (22) with the addition of FGF-coated beads from StemCultures. Pluripotency was evaluated by the ThermoFisher Scorecard (56), which compares gene expression of various self-renewal/pluripotency genes to that of genes distinct to the three germ layers (**Fig.S1A**). The pluripotency was also assessed in each cell line via immunofluorescence staining for several markers of pluripotency (**Fig. S1B**). Finally, karyotyping was performed to ensure that no genomic abnormalities had arisen during reprogramming (**Fig. S1C**). After gene editing, we used an allele-specific PCR screen to select for edited clones, which were verified by Sanger sequencing (**Fig. S2A**). Chromatograms from the target region are shown for the parental line as well as the corrected clones (**Fig. S2B**). We confirmed that there were no off-target mutations via Sanger sequencing of the top 20 off-target regions within exons (**Table S1**). *KLHL16* gene expression (**Fig. S2C**) was measured via sequence-specific primers (**Table S3**) and pluripotency and trilineage differentiation were assessed in the edited clones **(Fig. S2D-F)**.

### iPSC culture and differentiation

iPSCs were maintained on Matrigel (Corning, Cat. No. 354480) in StemFlex Medium (Thermo Fisher Scientific, Cat. No A3349401) and passaged every 3-4 days with 0.5 mM EDTA dissociation solution. Astrocytes were differentiated as described (22). To generate neural progenitor cells (NPCs), embryoid bodies (EBs) were plated on poly-ornithine and laminin coated plates in Neural Induction Medium (StemCell Technologies) and rosettes were selected after 12 days and expanded in Neural Progenitor Medium (StemCell Technologies). NPCs were differentiated to astrocytes using the Stem Cell Technologies STEMdiff astrocyte differentiation and maturation kits during which they were split weekly with Accutase (Millipore).

Organoids were generated using an established protocol (57) with a few following modifications for feeder-free conditions. The EBs were generated as described above on days 0-6. On day 7, the EBs were moved to neural medium (NM, Invitrogen) containing serum-free Neurobasal medium with B-27 without vitamin A (Invitrogen), GlutaMax (Gibco). NM was supplemented with FGF2 (20ng/mL, R&D Systems), and EGF (20ng/mL, R&D Systems). Media was changed daily for days 7-16 and every other day for days 17-25. After neural induction, the media was replaced with NM supplemented with BDNF (20ng/mL, Peprotech), and NT3 (20ng/mL, Peprotech) every other day to promote differentiation. On day 43, NM without growth factors was used for medium changes every four days.

### Preparation of protein lysates and immunoblotting

Total lysates were prepared from 60-80% confluent iPSCs plated on 6-well plates by rinsing cells quickly with 1 mL of 1X PBS and adding 2X Novex^™^ Tris-Glycine SDS Sample Buffer (ThermoFisher, Cat#LC2676) directly to the plate, collected into Eppendorf tubes, and heated at 95°C for 5 min. Protein lysates were normalized by running Coomassie staining before running western blots. For immunoblotting, samples were separated on a 4-20% gradient SDS-PAGE gel and transferred either for 1h at 110V or overnight at 40V onto an activated polyvinylidene difluoride membrane. Membranes were blocked in 5% non-fat milk dissolved into 0.1% Tween 20/PBS (PBST) for 1 hour at room temperature and then incubated in primary and secondary antibodies diluted into blocking solution at the concentrations listed above. Antibodies were detected via ECL reagents (PerkinElmer).

### RNA isolation and quantitative real-time PCR (qRT-PCR)

RNA was isolated from iPSCs using the ThermoFisher PureLink RNA mini kit (Thermo Fisher Scientific, Cat#12183025) following all manufacturer guidelines and immediately converted to cDNA. The High-Capacity cDNA Reverse Transcription kit (Thermo Fisher Scientific, Cat#4368814) was used to convert 2μg of total RNA to cDNA. To measure gene expression, qRT-PCR was performed using PowerUp SYBR Green Master Mix (Thermo Fisher Scientific, Cat#A25778) and the Applied Biosystems QuantStudio 6 Flex Real-Time PCR System. The primers used to detect gene expression in this study are listed in Table S2.

### *Fluorescent RNA in situ hybridization (ISH) to detect KLHL16* mRNA

Custom RNAScope® target-specific oligonucleotide (ZZ) probe (19ZZ length) was designed to target the region spanning nucleotides 2-997 of human *KLHL16* mRNA (NM_022041.4). The cells were treated with hydrogen peroxidase to minimize background (10 min, RT), followed by three PBST washes and permeabilization using Protease III digestion (1:15 dilution) (322000, ACD), at 40°C for 10 min. Subsequently, the expression of target probe was assessed using RNAscope® Multiplex Fluorescent v2 Assay protocol (323100, ACD). Positive (*POLR2A, PPIB*, and *UBC*) and negative (*dapB*) control probes were included to ensure specificity of signal detection. Following ISH signal development, the cells were washed three times in PBS for 5 minutes each and stained for vimentin as described below.

### Immunofluorescence, imaging, and analysis

Cells were fixed in methanol at −20°C for 10 min, washed three times in PBS, and blocked in Buffer B (2.5% Bovine Serum Albumin (Sigma), 2% normal goat serum (Gibco)/PBS) for 1 hour at room temperature. Next, cells were incubated in primary antibodies overnight at 4°C after which they were washed three times in PBS and incubated with Alexa Fluor-conjugated secondary antibodies 1 hr at room temperature. Cells were washed three times in PBS, incubated in DAPI (Invitrogen), washed twice in PBS, and mounted in Fluoromount-G (SouthernBiotech) overnight. Cells were imaged on Zeiss 880 confocal laser scanning microscope using a 40x oil immersion objective or for widefield on the EVOS-FL auto cell imaging system (Thermo Fisher Scientific) using a 20x (0.75 NA) objective. NIH ImageJ software was used to measure vimentin signal, size and intensity of the vimentin cytoskeleton as shown in Figure 4 by using the polygon tool to circle the vimentin signal and the measure tool, which provides a measure of area and average signal intensity. For analysis of GAN mRNA nuclear content in Figure 7, 3D Images were acquired using Zeiss confocal LSM880 microscope (Carl Zeiss) using a 63x oil immersion objective with NA 0.95 providing a 257nm resolution and using z-stack function at multiple areas. CellProfiler (v. 4.2.1) object identification pipelines were used to determine nuclear and perinuclear area, and to determine cells with and without vimentin aggregates (clumps) within a pre-determined perinuclear area. This was followed by identifying and counting GAN mRNA objects in nuclear area of cells with and without perinuclear clumps.

### Statistics

The graph data were presented using GraphPad Prism software and statistics were performed using the tests described in each figure legend. Adobe Photoshop was used to perform densitometry on immunoblots and Image J was used for signal quantification of immunofluorescence data.

## Acknowledgments

This study was funded by Hannah’s Hope Fund and NIH grants F99AG068523 and R21NS121578. We thank Dr. Anne Messer (Neural Stem Cell Institute) for feedback and advice on the studies and manuscript.

