## Supplemental Tables 1-3 for "Intermediate filament dysregulation and astrocytopathy in the human disease model of *KLHL16* mutation in giant axonal neuropathy (GAN)"

**Table S1: Summary of off-target Sanger sequencing from CRISPR/Cas9 editing of Patient 7**

| **Gene name** | **Gene ID** | **Parental** | **Clone B3** | **Clone G5** | **Clone 2D1** | **Clone 2D3** |
| --- | --- | --- | --- | --- | --- | --- |
| ATXN2L | 11273 | Homozygous: T>A (chr.16, 28832748) | Same as parental | Same as parental | Same as parental | Same as parental |
| BNIP3L | 665 | N.M. | N.M. | N.M. | N.M. | N.M. |
| CLRN1-AS1 | 116933 | N.M. | N.M. | N.M. | N.M. | N.M. |
| URI1 | 8725 | N.M. | N.M. | N.M. | N.M. | N.M. |
| SCN9A | 6335 | Heterozygous: A>G, G>A (chr.2, 166303519, 166303436) | Same as parental | Same as parental | Same as parental | Same as parental |
| COL19A1 | 1310 | N.M. | N.M. | N.M. | N.M. | N.M. |
| RFLNB | 359845 | N.M. | N.M. | N.M. | N.M. | N.M. |
| CSN1S2AP | 286828 | N.M. | N.M. | N.M. | N.M. | N.M. |
| PDZD2 | 23037 | N.M. | N.M. | N.M. | N.M. | N.M. |
| PNLIPRP2 | 5408 | Heterozygous: G>C (chr. 10, 116624046) | Same as parental | Same as parental | Same as parental | Same as parental |
| DIP2A | 23181 | N.M. | N.M. | N.M. | N.M. | N.M. |
| EGFEM1P | 93556 | N.M. | N.M. | N.M. | N.M. | N.M. |
| GINS2 | 51659 | N.M. | N.M. | N.M. | N.M. | N.M. |
| SCN3A | 6328 | N.M. | N.M. | N.M. | N.M. | N.M. |
| SGMS1 | 259230 | N.M. | N.M. | N.M. | N.M. | N.M. |
| RUBCNL | 80183 | N.M. | N.M. | N.M. | N.M. | N.M. |
| c11orf58 | 10944 | N.M. | N.M. | N.M. | N.M. | N.M. |
| BTN1A1 | 696 | N.M. | N.M. | N.M. | N.M. | N.M. |
| USP50 | 373509 | N.M. | N.M. | N.M. | N.M. | N.M. |
| IRF4 | 3662 | N.M. | N.M. | N.M. | N.M. | N.M. |

*N.M. = no mutations

**Table S2: GAN (*KLHL16*) qRT-PCR primers**

| **Primer Name** | **Target** | **Sequence (5’** 🡪 **3’)** |
| --- | --- | --- |
| GAN_exon2-3_F | *GAN* | TCGGTAATGGTTATGAGAGAGATCC |
| GAN_exon2-3_R | *GAN* | TAAGGTCCGTCAGTAGCAGC |
| GAN_exon10-11_F | *GAN* | CCGCCAGTTCCTCTTTTGTT |
| GAN_exon10-11_R | *GAN* | GTCGGATGGAAGGAGTGGTTT |

**Table S3: *KLHL* gene family primers**

| **Primer Name** | **5' --> 3' Sequence** |
| --- | --- |
| hsKLHL1_F | AAACTCTTCAGCCACCCGTC |
| hsKLHL1_R | AGAAAGTGCTCACACCGCTT |
| hsKLHL2_F | CTGCCAACAGAGCAGCGTAT |
| hsKLHL2_R | CCGGATAGCCTTTGGTGCTT |
| hsKLHL3_F | GGAGGGTGAAAGTGTCAAGC |
| hsKLHL3_R | AACATCGCACAGAAGTAGGGG |
| hsKLHL4_F | TGCCCTCTTGGAGACAAACTC |
| hsKLHL4_R | AGGGTAGCTTCACCACTACA |
| hsKLHL5_F | GTGGCTGTACTGGAAGGTCC |
| hsKLHL5_R | GCCTGAGGGTCCCATCTTTC |
| hsKLHL6_F | CTCTTCCAGTTCCTGCGGAT |
| hsKLHL6_R | ACGGGCAGGTCAAGAAACTC |
| hsKLHL7_F | GAAACTGGGTCCTCCGACAC |
| hsKLHL7_R | CATTGGCTTCCACCATCCCA |
| hsKLHL8_F | CTGAAGCCAAGCAAACGCT |
| hsKLHL8_R | CAACAAGCTCTAGCCACCAGT |
| hsKLHL9_F | GCAGGAACCACACGCTTTTT |
| hsKLHL9_R | TTCCACCTGTGAACATGGCT |
| hsKLHL10_F | GTGCCTATCACACCGGACAA |
| hsKLHL10_R | ACCCCTGACGATACCCATGA |
| hsKLHL11_F | GCCTACCGATATTGTGCGGA |
| hsKLHL11_R | CGTGACAAGCAGTGGCTCTA |
| hsKLHL12_F | GTGGAGCTTTTTGCCAAGCA |
| hsKLHL12_R | ACTGAACTAAGGCGGGAACG |
| hsKLHL13_F | ATGGTGTGAGCAAAGTCGGT |
| hsKLHL13_R | TAGGTGTTGGCAATCCGTCC |
| hsKLHL14_F | TTGGGTGGAGAGGACCAGTG |
| hsKLHL14_R | TTTCCTGCATGGGTGGAAGTT |
| hsKLHL15_F | CGTGTTTGACCCCAGCAAAG |
| hsKLHL15_R | CACAGACACCACCGAAGACA |
| hsKLHL17_F | CCTCCAACTGCCTGGGTATC |
| hsKLHL17_R | TCCAGAACCTGTTTCAGGGGC |
| hsKLHL18_F | ATGTGTGCTGTGCTGTACGA |
| hsKLHL18_R | CAGGGCCAGGAACTCTTCTG |
| hsKeap1_F | TCTTCAACCTGTCCCACTGC |
| hsKeap1_R | TTCGCAGTCGTACTTGACCC |
| hsKLHL20_F | AAGCACCCTCGACAAACCTT |
| hsKLHL20_R | GACGGCTCTCTGCCAATTCT |
| hsKLHL21_F | CTATGACTGCGTGTGGAGGT |
| hsKLHL21_R | GGAGTCAGTGGTGTGGTCAT |
| hsKLHL22_F | TTTGTGGCCTTCTCTCGGAC |
| hsKLHL22_R | AACCGCACTGTCTCAAGGAG |
| hsKLHL23_F | GGGGCTACAGGACGGATAAC |
| hsKLHL23_R | CTTCTGCTGGAGCCCCTTTT |
| hsKLHL24_F | CTATGTTGCCGGTGGACTGA |
| hsKLHL24_R | CATTGCAGCTACCCCTGTGA |
| hsKLHL25_F | TTGCCTACTCCTCACGCATC |
| hsKLHL25_R | TGCGCCAGGAGAACTCATAC |
| hsKLHL26_F | CTCCTCGATGTTGTGCTGAC |
| hsKLHL26_R | CTGGGGAACAGTAGCCTGAAG |
| hsIPP_F | CTTTGGGTGGATGGGTTGGA |
| hsIPP_R | CCTTGCATTTCACAGCACCC |
| hsKLHL28_F | TTCCTACCCTGCTCTGTACC |
| hsKLHL28_R | CAGAGTTCGTGATGTTGGCG |
| hsKLHL29_F | TGGAGTTTGTCTACACGGGC |
| hsKLHL29_R | CGAGAAAGGACACGCAGACT |
| hsKLHL30_F | CCATGTTTGCGGGTGACTTC |
| hsKLHL30_R | ACGAAGTCCACCAGTTGTCC |
| hsKLHL31_F | AGGCGAACAAGAATCCGAGG |
| hsKLHL31_R | CCGTAAGCTTGCTCCATCCA |
| hsKLHL32_F | AAGTGGATAAGCCGTAGCCC |
| hsKLHL32_R | CATTTTGGCCAGTCAGCAGG |
| hsKLHL33_F | TTTGGAGGGCCAGTTGTACG |
| hsKLHL33_R | TCAGAAACGTCCCTGGCTTC |
| hsKLHL34_F | GGAGCAGGTTTGGAGCAAGA |
| hsKLHL34_R | CAGAGAAGGGCTCGTATCGC |
| hsKLHL35_F | GGCAAGCTCTTCGTGATTGG |
| hsKLHL35_R | TCCTTGGGGTCAAAGCACTG |
| hsKLHL36_F | TGCGGCCTCCAATCTTCTTT |
| hsKLHL36_R | AGGTAGAAATCCACACGGCG |
| hsENC1_F | GTGGACCAAGGTGGGAGATG |
| hsENC1_R | GCATCGCTGAATGCCAAAGT |
| hsKLHL38_F | AGAACCCTGTGCGCCTTATC |
| hsKLHL38_R | ATGACAATCCGCTCCCCAAG |
| hsIVNS1ABP_F | AATCAACTGGGTGCAGCGTA |
| hsIVNS1ABP_R | AACACCTCAGCCTGTCCATC |
| hsKLHL40_F | GAGAGATCAAGGACGGCGAG |
| hsKLHL40_R | GGGTCCGATTCACCCCATTT |
| hsKLHL41_F | CTTCTTGACTGCCCGAGACT |
| hsKLHL41_R | CGCACCCATTTCATCACTGC |
| hsKLHL42_F | TTACAACCCCGAGCAGGATG |
| hsKLHL42_R | GTTCATGTTCCGGTCTCTGGT |
