## Supplementary figures and images for "Intermediate filament dysregulation and astrocytopathy in the human disease model of *KLHL16* mutation in giant axonal neuropathy (GAN)"

### Supplemental Figure 1

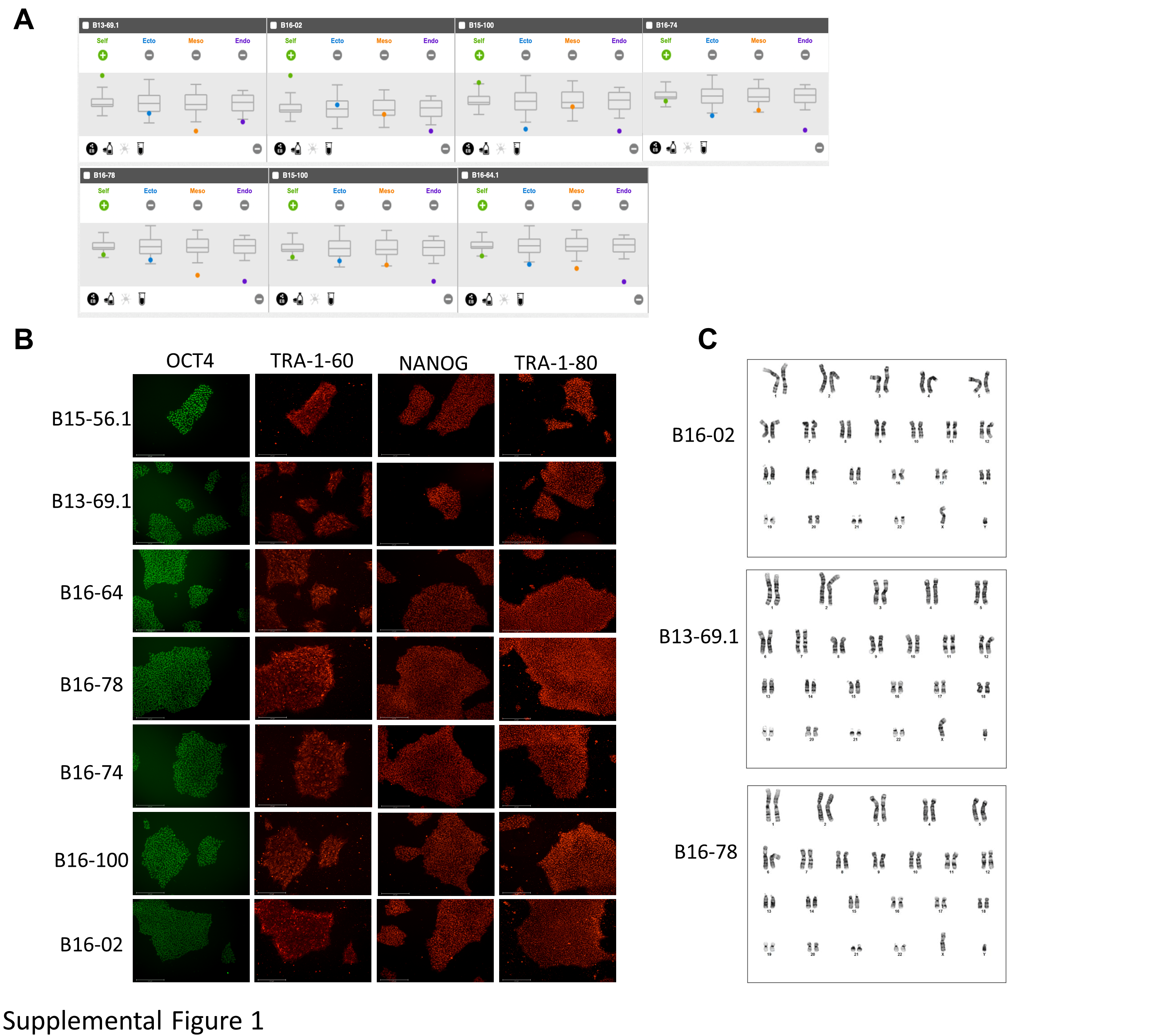

### Supplemental Figure 2

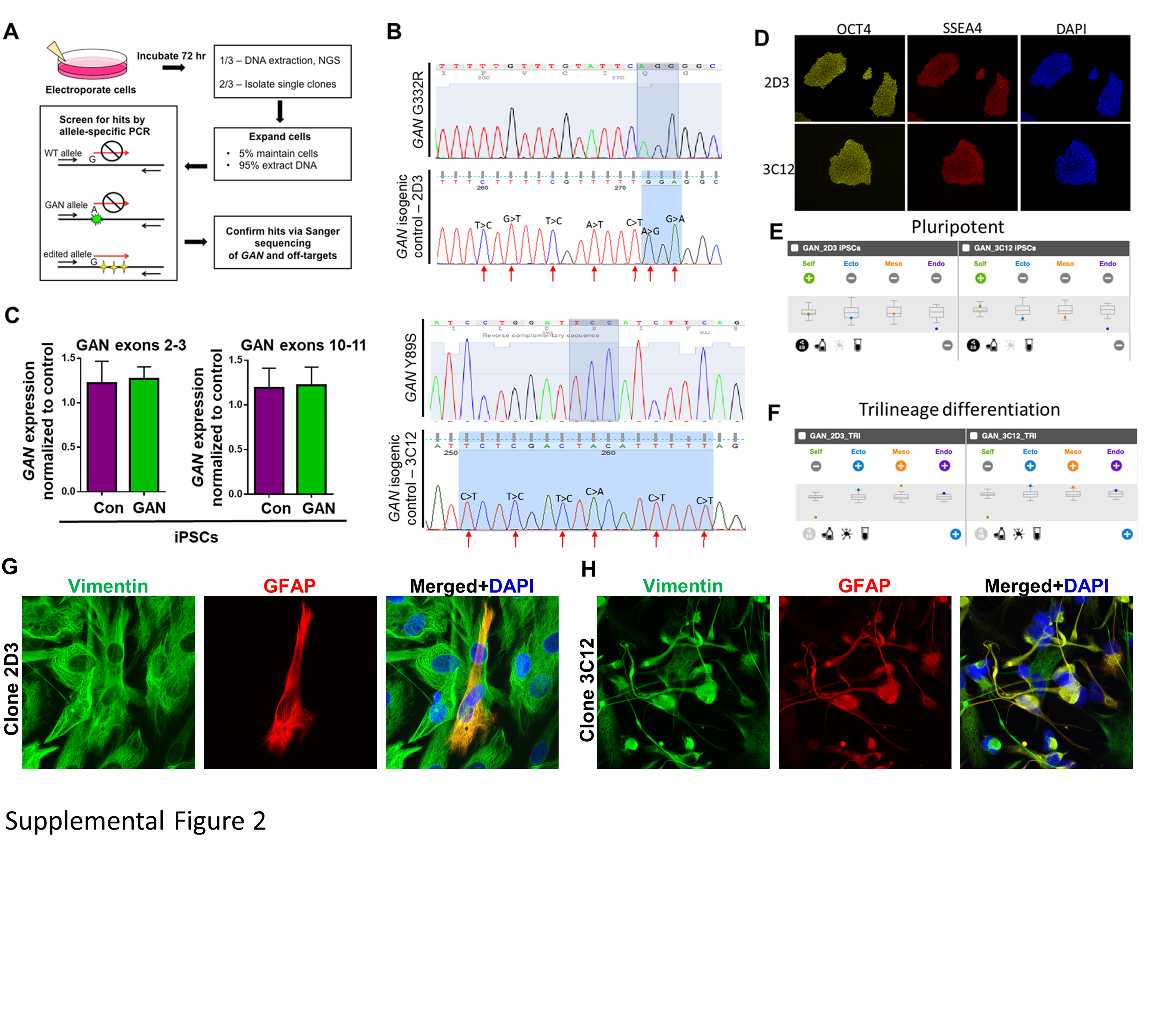

### Supplemental Figure 3

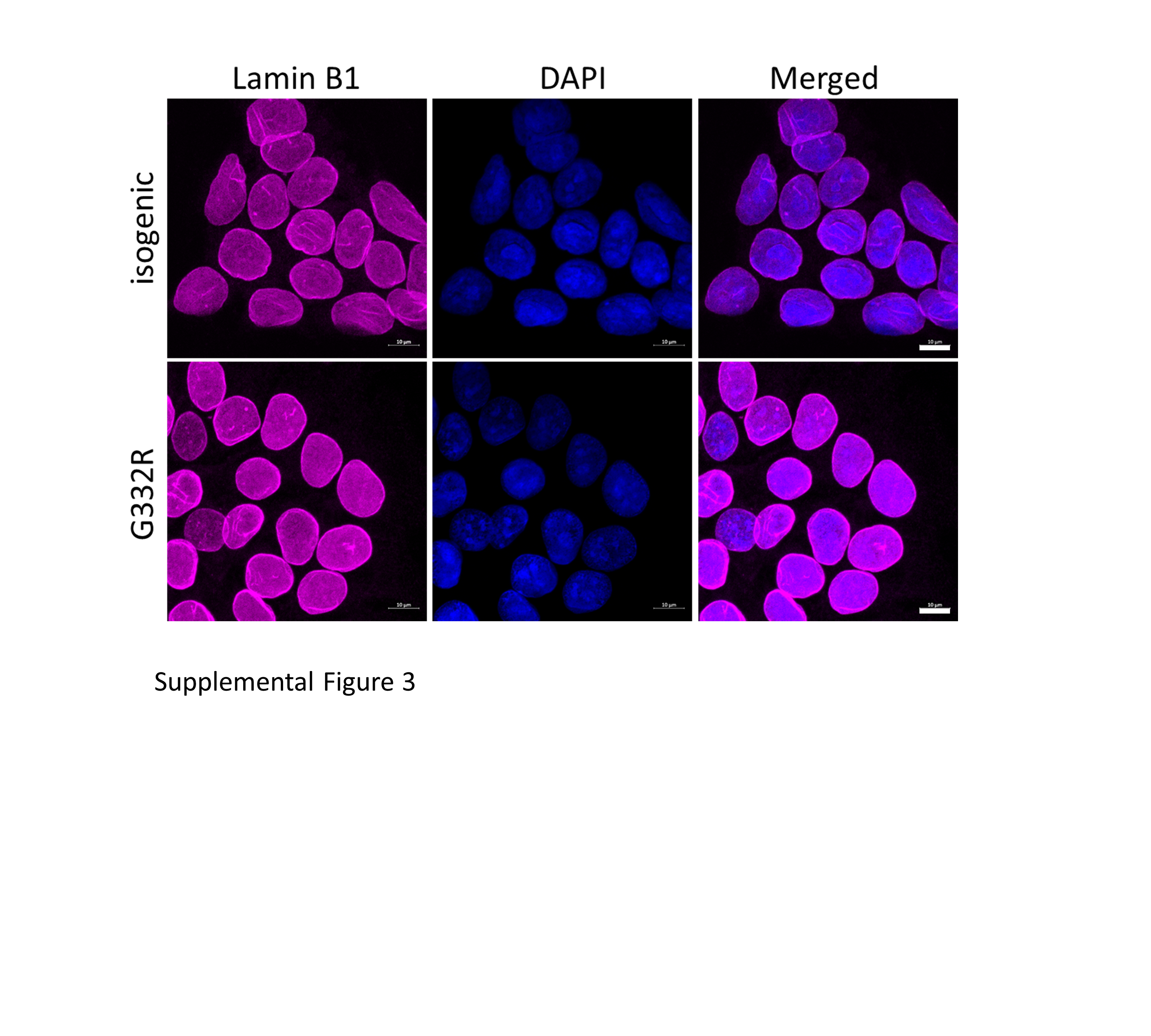

### Supplemental Figure 4

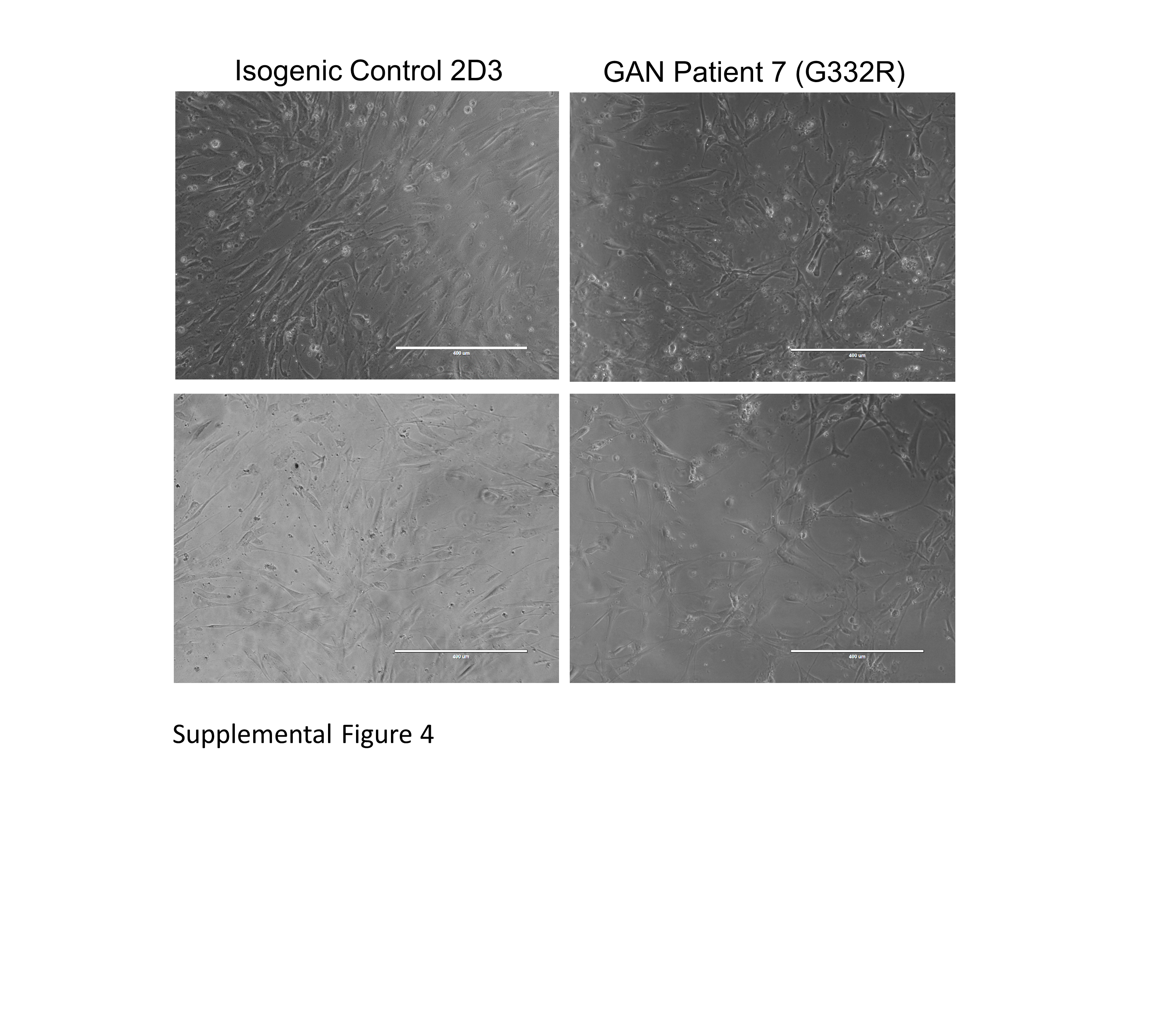

### Supplemental Figure 5

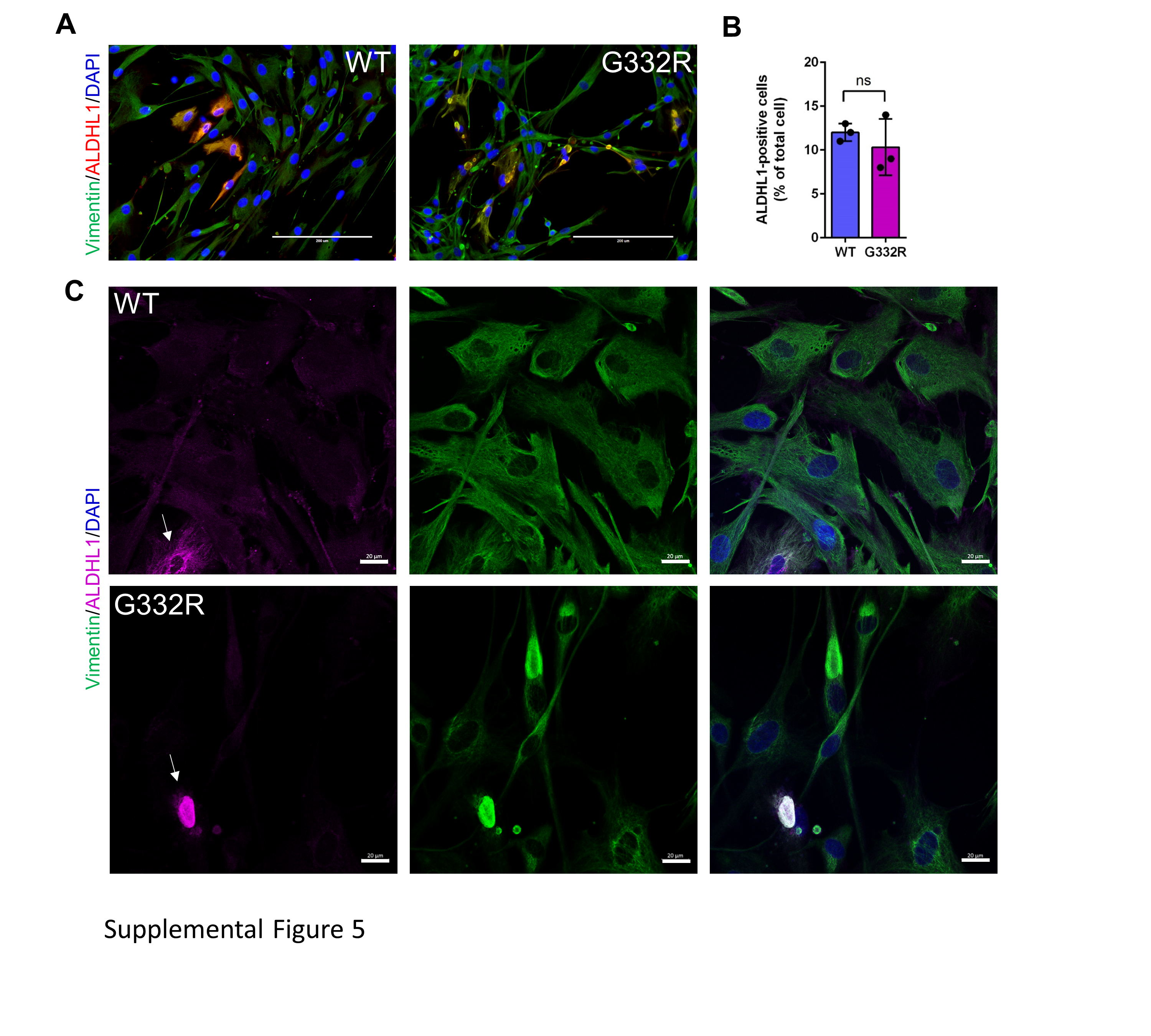

### Supplemental Figure 6

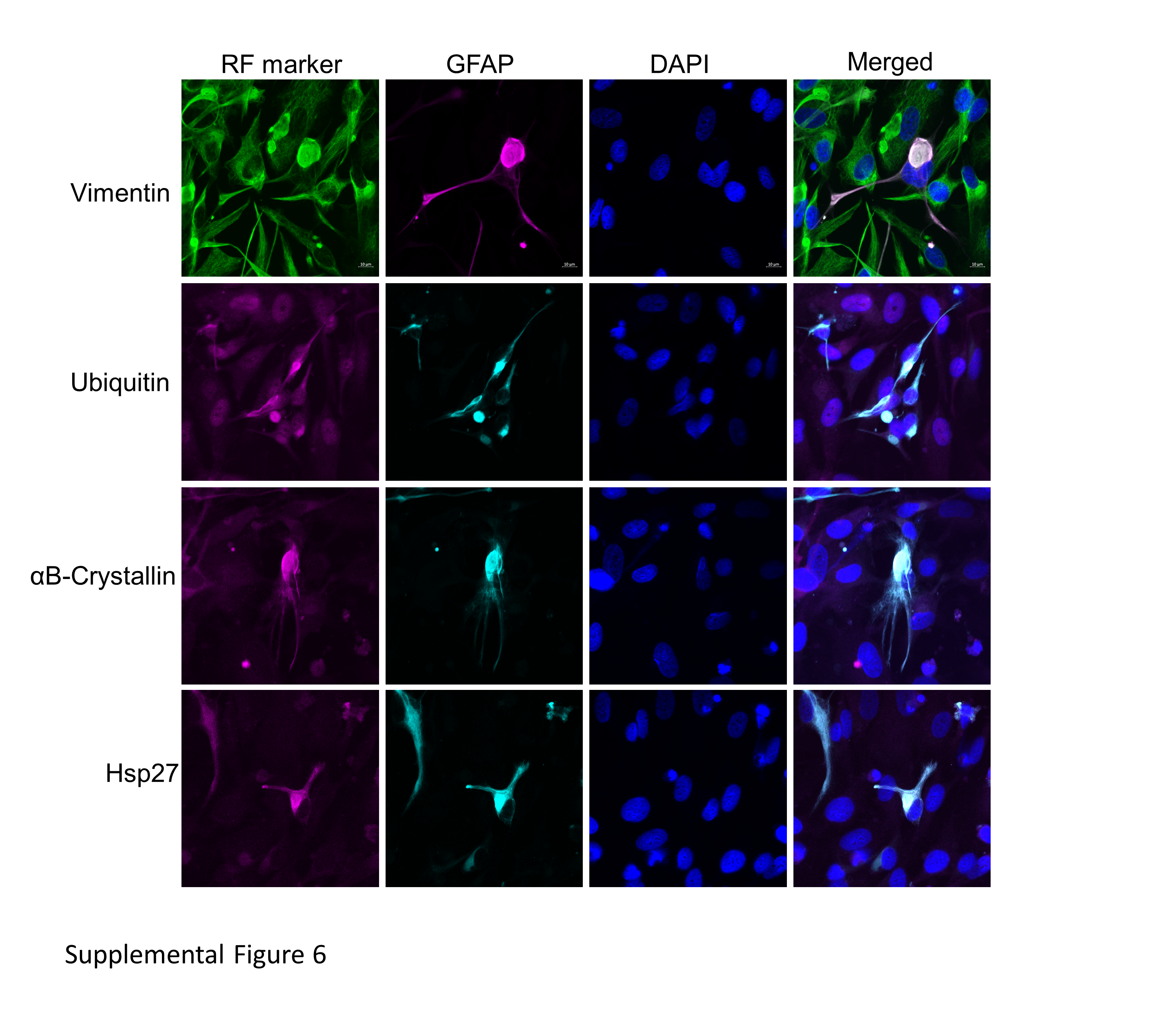
