## Supplemental Figure Legends for "Intermediate filament dysregulation and astrocytopathy in the human disease model of *KLHL16* mutation in giant axonal neuropathy (GAN)"

**Battaglia et al. BioRxiv 2023**

**Supplemental Figure Legends**

**Supplemental Figure 1.** Generation of *GAN* iPSCs from patient fibroblasts. **(A-B)** Pluripotency was assessed via ThermoFisher Taqman scorecard analysis and immunofluorescence staining for OCT4 (green), TRA-1-60, NANOG, and TRA-1-80 (red) pluripotency markers. **(C)** GAN iPSCs were characterized for genomic stability via karyotyping as shown by representative images from GAN patients 2, 4, and 7 (please also see Figure 1C).

**Supplemental Figure 2.** Correction of *KLHL16* mutations via CRISPR/Cas9 gene editing. **(A)** Gene editing strategy for GAN Patient 7 iPSCs. The G is the wild-type allele, the green shape represents the GAN mutation (G>A), and the yellow stars represent the silent mutations introduced by the repair construct. B lack arrows depict universal primers, whereas the red arrow is an allele specific primer that exclusively binds to the corrected sequence. **(B)** Chromatograms display the original GAN mutant sequence (G332R, GGG>AGG; Y89S, TAC>TCC) and the sequence from a corrected clone where silent mutations are indicated by red arrows. **(C)** *GAN* gene expression in iPSCs by qRT-PCR using primers at the beginning (left) and end (right) of the gene. **(D)** Immunofluorescence staining for pluripotency markers OCT4 and SSEA4 expression in isogenic control lines 2D3 and 3C12. Pluripotency **(E)** and trilineage differentiation **(F)** were assessed in clone 2D3 via ThermoFisher Taqman scorecard analysis. **(G)** Immunofluorescence analysis of IF proteins vimentin (green) and GFAP (red) in isogenic clone 2D3 iPSC-astrocytes from patient 7 (correction of G332R mutation). **(H)** Immunofluorescence analysis of IF proteins vimentin (green) and GFAP (red) in isogenic clone 3C12 iPSC-astrocytes from patient 2 (correction of the Y89S mutation).

**Supplemental Figure 3**. Confocal imaging of lamin B1 (magenta) and DAPI (blue) in GAN patient 7 (G332R) and corresponding isogenic control iPSCs (clone 2D3). Scale bars=10um.

**Supplemental Figure 4.** Bright field images of isogenic control 2D3 (left panels) and parental line from GAN patient 7 (G332R) iPSC-astrocytes at day 60.

**Supplemental Figure 5. ALDHL1 expression in GAN iPSC-astrocytes. (A)** GAN patient 7 (G332R) and corresponding isogenic control (WT; clone 2D3) iPSC-astrocytes were stained for ALDHL1 (red), vimentin (green), and DAPI (blue) Scale bars=200 μm. **(B)** Quantification of ALDHL1-positive cells as a percentage of total cells from 3 representative fields of view (n=136 WT; n=216 G332R total cells counted). n.s.=not significant; unpaired t-test. **(C)** GAN patient 7 (G332R) and isogenic control iPSC-astrocytes (WT; clone 2D3) were stained for ALDHL1 (magenta), vimentin (green) and DAPI (blue). Scale bars=10 μm. Arrows highlight ALDHL1-positive cells in the representative images.

**Supplemental Figure 6.** Expression of IF proteins vimentin and GFAP and Rosenthal Fiber markers in GAN iPSC-astrocytes. Images show co-staining for vimentin (green) and GFAP (magenta) in the top row. The bottom three rows show co-staining for Rosenthal Fiber markers ubiquitin, αB-crystallin and Hsp27 (magenta) with GFAP (cyan) and DAPI (blue). Patient 2 (Y89S mutation) parental line GAN iPSC-astrocytes were used in this analysis. Scale bar=10μm.
